# BioFrame: Enhancing Reproducibility and Accessibility in Genomics through Web-Based Workflow Design

**DOI:** 10.1101/2025.10.11.681789

**Authors:** Mohamed Salim Nasser Al Hinai, Zakira Naureen, Syed Abdullah Gilani

## Abstract

Since, Bioinformatic research is getting attraction that can be seen with sudden increase in development of tools as well as publications. However, there are challenges in bioinformatics which are making the results difficult to obtain while facing reproducibility, scalability, and accessibility of computational workflows. Though command line tools are powerful but majority of the bioinformatic researchers have little or no expertise in the extensive computational skills.

To address these issues, we present here the easy to use and getting results with ease through our tool, BioFrame. BioFrame employs a containerized microservices architecture orchestrated via Docker Compose, separating concerns across presentation, orchestration, execution, and storage layers. The bioinformatic tools are encapsulated here as versioned container images with declarative metadata schemas, enabling dynamic tool discovery and parameter validation. Therefore, through user designed workflows one can easily get directed acyclic graphs while using the web interface.

## Introduction

In the current era of genomics advancements, the genome analysts need to use the complicated pipelines with ease especially those who do not have command on Linux systems as well as Python or know how about commands used in Linux related software for genomes. Numerous workflow management systems (WMSs) have been developed to date. Some, such as Galaxy and KNIME, provide user-friendly graphical interfaces and do not require command-line interaction. Others are based on domain-specific languages (DSLs), including Nextflow (Di Tommaso et al., 2017), Snakemake (Köster *et al*., 2012), GenPipes (Bourgey *et al*., 2019), bPipe, and Pachyderm (Novella *et al*., 2019), allowing users to define workflows using custom syntax. Tools like SciPipe and Luigi are library-based and require programming skills to implement workflows. Additionally, systems such as Cromwell (using WDL), cwltool (with CWL), and Toil (which supports CWL, WDL, and Python) combine both workflow specification and execution capabilities, offering flexibility and standardization for complex analyses (Wratten et al., 2021). The complexity of existing bioinformatics pipeline frameworks presents significant barriers for researchers without strong computational backgrounds. These challenges not only limit accessibility but also compromise reproducibility when workflows cannot be easily shared or validated.

A critical step in modern genomic analysis involves processing raw sequencing reads—whether single-end or paired-end—through a series of tasks such as quality assessment using tools like FastQC, trimming through different tools such as Trimmomatic (Bolger *et al*., 2014), genome assembly with SPAdes (Bankevich *et al*., 2012; Prjibelski *et al*., 2020), scaffold generation, annotation, and alignment with reference or publicly available genomes **(Table 1)**. A wide range of trimming tools are available, including Trimmomatic (Bolger et al., 2014), Cutadapt (Martin, 2011), Fastp (Chen et al., 2018), Trim Galore, BBDuk (BBTools suite), Sickle, AfterQC (Chen et al., 2017), Skewer (Jiang et al., 2014), and NGS QC Toolkit (Patel & Jain, 2012). These steps are often technically demanding, time-consuming, and require high-performance computing environments typically available only on Linux systems. Standard Windows machines often lack the necessary memory, storage, and command-line support to run these tools efficiently, especially in graphical modes.

One of the major challenges in bioinformatics is dependencies on managing software, where tools often have complex and conflicting requirements. The bioinformatic tools such as **Anaconda** bundles the **Conda** package and environment manager, enabling users to install and manage software in isolated environments across different operating systems (Anaconda, 2024). **Bioconda**, a community-maintained Conda channel, curates thousands of bioinformatics packages and interoperates with container registries such that Bioconda packages are automatically available as Docker and Singularity images through the BioContainers project (Grüning et al., 2018). In contrast, **Docker** and **Singularity** offer a more comprehensive level of encapsulation by bundling the entire execution environment— including the OS, dependencies, and software—into portable, versioned container images that guarantee reproducibility and portability across varied computational environments (Merkel, 2014; Kurtzer et al., 2017). While Conda (and by extension Anaconda and Bioconda) simplifies environment and package management, containerization provides stronger isolation, consistency, and ease of sharing for complex bioinformatics workflows.

In recent years, containerization has emerged as a powerful paradigm in bioinformatics for improving the reproducibility, portability, and scalability of computational workflows. Containers, such as those built with Docker or Singularity, package software together with all its dependencies, ensuring consistent behavior across different computing environments. This is particularly beneficial in genomics, where diverse tools and libraries are often required within a single pipeline.

Several modern workflow management systems (WMSs) have integrated containerization natively. Nextflow (https://github.com/nextflow-io/nextflow), Snakemake (https://snakemake.github.io/), Cromwell (https://github.com/broadinstitute/cromwell), Toil, Pachyderm, and Galaxy support container-based execution using Docker or Singularity, enabling workflows to be executed seamlessly across local machines, HPC clusters, or cloud environments. These tools leverage containerization to ensure that bioinformatics pipelines are reproducible and easier to deploy.

In contrast, older or simpler systems such as bPipe, Luigi, and GenPipes do not natively depend on containers and often require manual installation and configuration of software dependencies on the host system (Sadedin *et al*., 2012). This can lead to reproducibility issues and limits portability, particularly in collaborative or heterogeneous computing environments.

To address these challenges, we developed BioFrame, a web-based platform that abstracts the complexity of command-line tools and high-performance computing, enabling users to perform comprehensive genomic analyses through an intuitive, reproducible, and scalable interface without requiring extensive computational expertise.

By adopting a container-based architecture, platforms like BioFrame further reduce technical barriers for researchers, allowing users to execute complex genomic workflows through a web interface without needing to manage software installations or system configurations manually.

## Materials and Methods

### SYSTEM DESIGN AND ARCHITECTURE

#### Overall System Architecture

BioFrame was designed following a three-tier architecture pattern to ensure modularity, scalability, and maintainability. This architectural approach separates concerns into distinct layers, each with well-defined responsibilities and interfaces. The system comprises three principal components: the presentation layer, the orchestration layer, and the containerization layer **(Figure 1)**.

**Figure 1:**
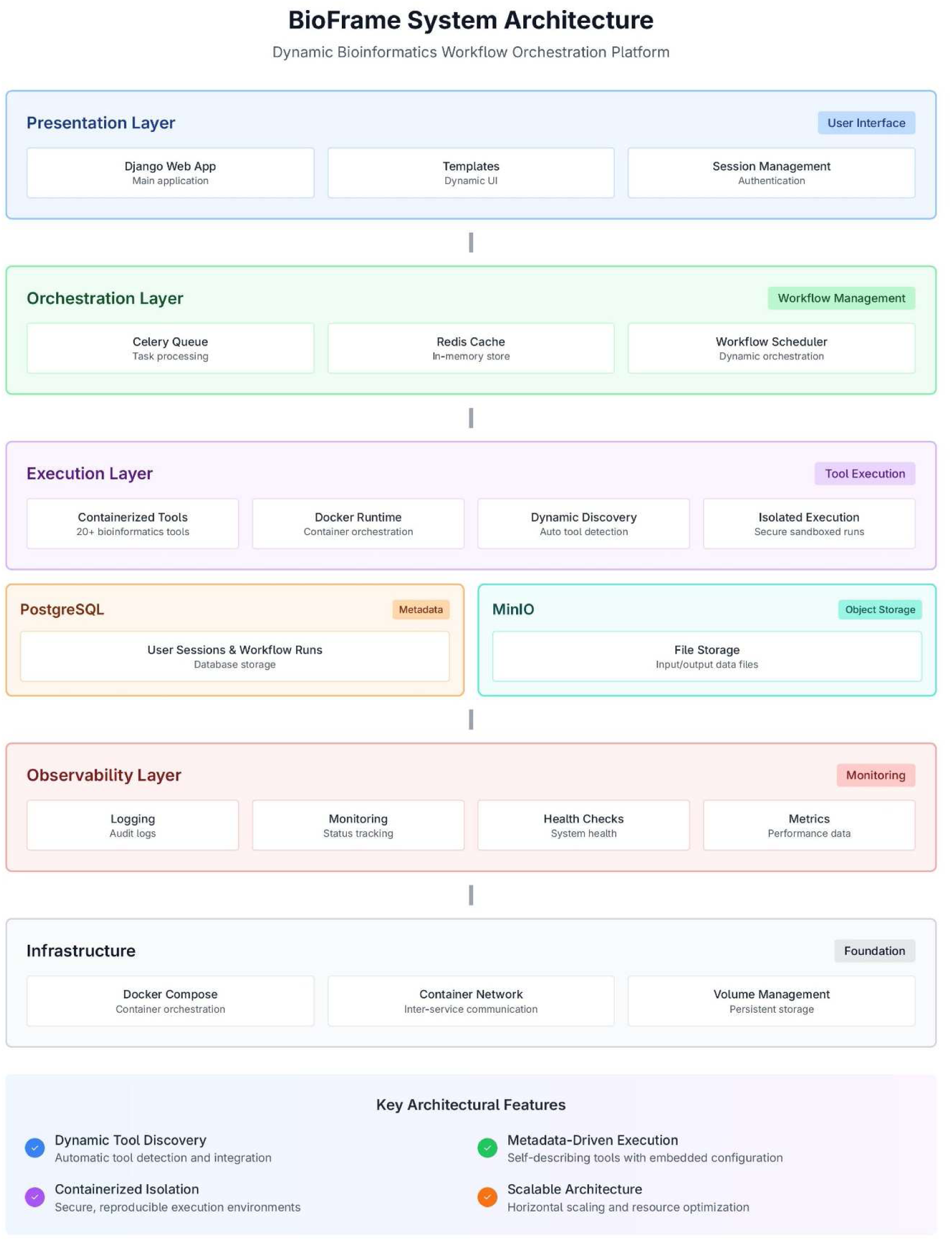
BioFrame System Architecture.

##### Presentation Layer (Portal)

A Django-based web application (Python 3.8+) serves as the primary user interface, implementing the Model-View-Template (MVT) architectural pattern. This layer handles *user authentication*, *workflow configuration*, *real-time monitoring*, and *result visualization*. The portal communicates with the orchestration layer through a RESTful API interface and manages user sessions using session-based authentication with CSRF protection. The interface was designed following principles of user-centered design, with particular attention to accessibility for bioinformatics researchers who may have limited command-line experience.

##### Orchestration Layer (Workflow Engine)

The core workflow orchestrator, implemented in Python, manages the entire lifecycle of bioinformatics pipelines from initialization through completion. This layer implements a metadata-driven execution model that dynamically discovers and integrates containerized tools without requiring code modifications. The orchestrator performs directed acyclic graph (DAG) validation to ensure work-flow structural integrity, topological sorting for execution order determination, and container lifecycle management through the Docker Engine API. Resource allocation, error detection, and recovery mechanisms are all handled within this layer.

##### Containerization Layer (Tools)

Each bioinformatics tool is packaged as an isolated Docker container with embedded metadata specifications. This layer provides **reproducible execution environments** with precise dependency management, resource allocation, and cross-platform compatibility. Tools are completely self-contained, including all required libraries, dependencies, and configuration files. The containerization approach ensures that workflows produce identical results regardless of the host system configuration, addressing a critical challenge in computational biology research **(Figure 2)**.

**Figure 2:**
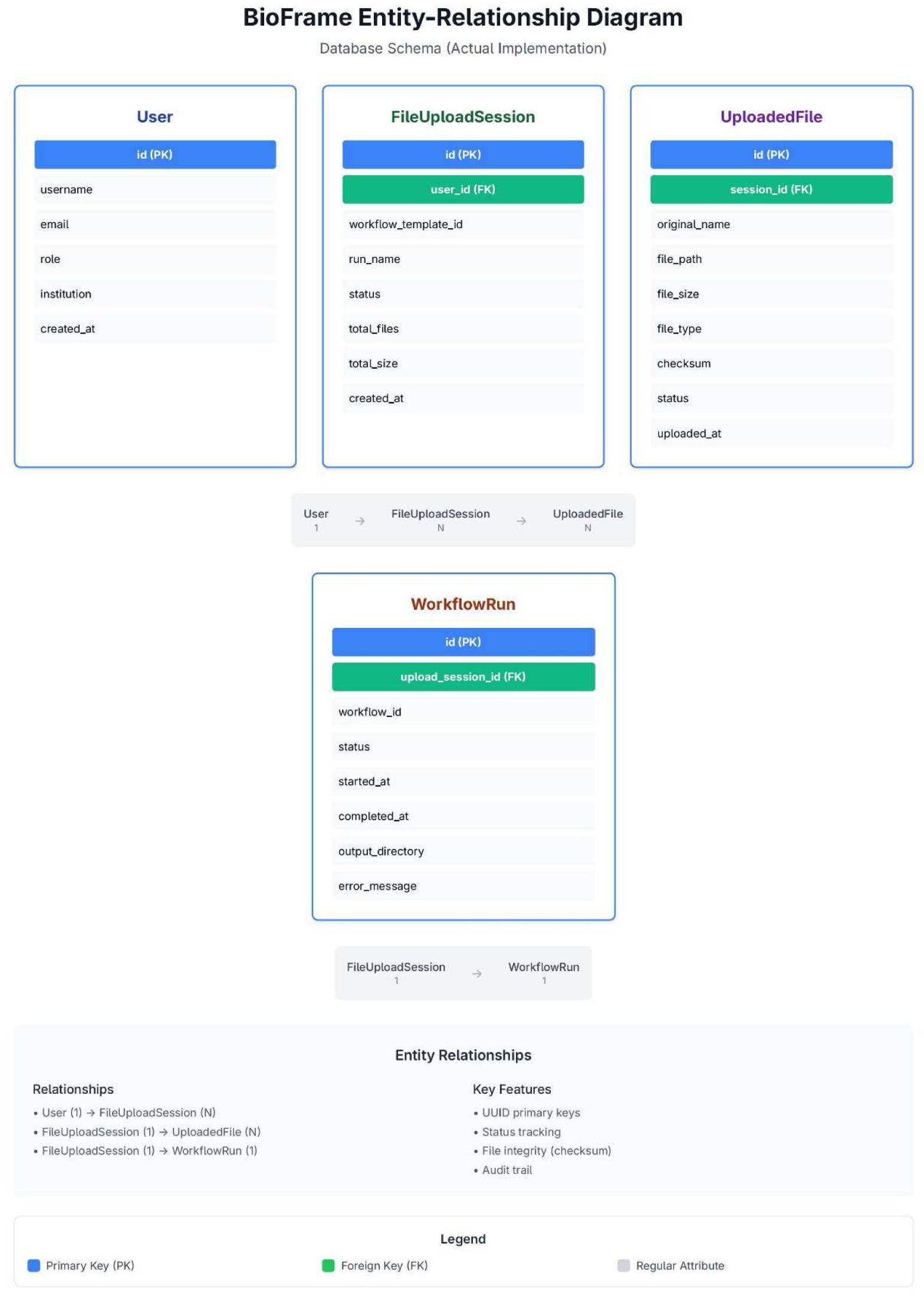
BioFrame Entity Relationship Diagram.

#### Data Model Design

The data model was designed to support complex workflow orchestration while maintaining referential integrity and query performance. Following principles of normalized database design, the entity-relationship diagram (ERD)comprises eight core entities with carefully defined relationships **(Figure 3)**.

**Figure 3:**
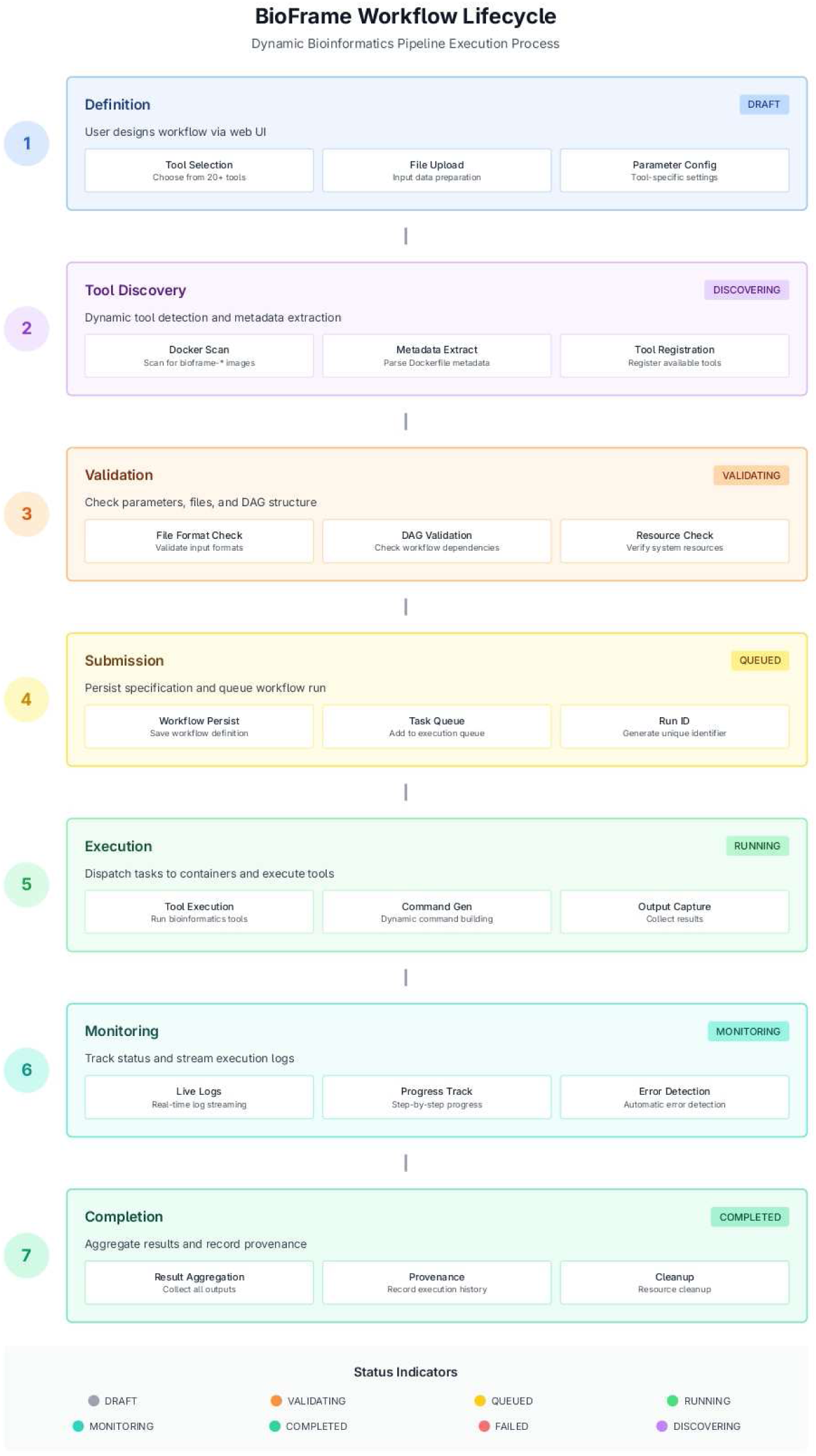
BioFrame Workflow Lifecycle.

##### Core Entities

The **Tool** entity represents bioinformatics software packages with attributes including name, category, and description. Each tool has a one-to-many relationship with **Version** entities, which represent specific releases with version numbers, container images, and release dates. The **Parameter** entity defines input specifications with type validation, default values, and requirement flags, maintaining a many-to-one relationship with versions.

The **Workflow** entity represents directed acyclic graphs of tools with metadata including name, description, creation date, and owner. Workflows maintain many-to-many relationships with tool versions through inter-mediate step entities, and many-to-one relationships with users. **Run** entities represent execution instances with status tracking, timestamps, and execution metadata, while **Task** entities represent individual tool executions within workflow runs with detailed logging.

The **File** entity captures input/output data artifacts with checksums (SHA-256), size, and type metadata, maintaining many-to-many relationships with tasks for tracking data lineage. Finally, the **User** entity manages authenticated researchers with role-based access control, institutional affiliations, and usage quotas.

##### Database Implementation

The database schema was implemented using PostgreSQL 15 with Django ORM for object-relational mapping, ensuring ACID compliance and transaction management. Indexes were strategically placed on frequently queried fields (work flow_id, user_id, status, timestamp) to optimize query performance **(Table 8)**. Database query response times averaged less than 100ms for metadata queries under normal operating conditions.

#### Containerization Strategy

Each bioinformatics tool was containerized using Docker with Ubuntu 22.04 LTS as the base image to ensure stability and broad package availability. The containerization strategy followed industry best practices for reproducibility, security, and performance.

##### Base Image Selection

Ubuntu 22.04 LTS was chosen for its five-year support lifecycle, extensive package ecosystem, and widespread adoption in the bioinformatics community. Alternative base images (Alpine Linux, CentOS) were evaluated but rejected due to compatibility issues with certain bioinformatics tools requiring GNU libc rather than musl libc.

##### Dependency Management

All dependencies were explicitly version-pinned to ensure reproducibility. For example, rather than installing “bwa” generically, installations specified exact versions (e.g., “bwa=0.7.17-r1188”). Dependency resolution was handled through apt-get for system packages and conda / pip for language-specific dependencies, with version constraints documented in each Docker file.

##### Directory Structure

A standardized directory structure was enforced across all containers: /data for in-put files, /output for results, and /logs for execution logs. This consistency simplified volume mounting and path resolution during workflow execution. Working directories were set to /data by default to ensure tools could locate input files without explicit path specifications.

##### Security Considerations

Where possible, containers executed as non-root users to minimize security risks. User ID mapping ensured that output files were owned by the appropriate user on the host system. Container capabilities were limited using Docker’s security features, preventing privilege escalation and reducing the attack surface.

## Results and Discussion

### DYNAMIC TOOL INTEGRATION SYSTEM

#### Metadata Specification Framework

A critical innovation of BioFrame is the metadata-driven tool integration system. Rather than requiring manualcode modifications for each new tool, the system extracts configuration information directly from structured meta-data embedded in tool Dockerfiles. This approach represents a paradigm shift from traditional static tool integra-tion to dynamic, self-describing tools.

Each tool’s Dockerfile includes a structured metadata block beginning with the commentBIOFRAME_TOOL_METADATA. This block contains 24 standardized fields organized into six categories: coreidentification, input/output specifications, execution configuration, quality assurance, and user interface elements.

##### Core Identification Metadata

The tool_name field specifies the display name presented to users in theweb interface. The tool_description provides a concise functional description explaining the tool’s purposeand capabilities. The tool_version field uses semantic versioning (MAJOR.MINOR.PATCH) to track tool re-leases. The tool_category field enables taxonomic classification (e.g., Quality Control, Assembly, Alignment,Variant Calling) for organizing tools in the user interface. Additional fields include tool_author for attributionand tool_url linking to official documentation.

##### Input/Output Specifications

The tool_input_formats field defines a comma-separated list of ac-cepted file formats (FASTQ, FASTA, BAM, VCF, etc.), enabling automatic validation of workflow connections.The tool_output_formats field specifies expected output formats, allowing the system to verify that consec-utive tools in a workflow are compatible. The tool_expected_outputs field lists specific file paths or pat-terns that should exist after successful execution, supporting output validation.

##### Execution Configuration

The tool_commands field enumerates available executable commandswithin the container. The tool_primary_command identifies the main executable for the tool. Most impor-tantly, the tool_command_template field provides a parameterized command template with placeholders thatare substituted at runtime. Resource requirements are specified through tool_memory_requirement (e.g.,“4g”, “8g”) and tool_cpu_requirement (number of cores), enabling resource-aware scheduling **(Table 7)**.

##### Quality Assurance

The tool_success_indicators field contains regex patterns or literal stringsthat, when found in execution logs, indicate successful completion. Conversely, tool_failure_indicatorsdefines patterns signaling errors or failures. This mechanism enables tool-specific error detection far more precisethan relying solely on exit codes.

#### Dynamic Tool Discovery Algorithm

The orchestrator implements an automatic tool discovery algorithm that executes during system initialization,eliminating the need for maintaining hardcoded tool registries. This algorithm represents a key contribution to ex-tensibility and maintainability.

##### ALGORITHM

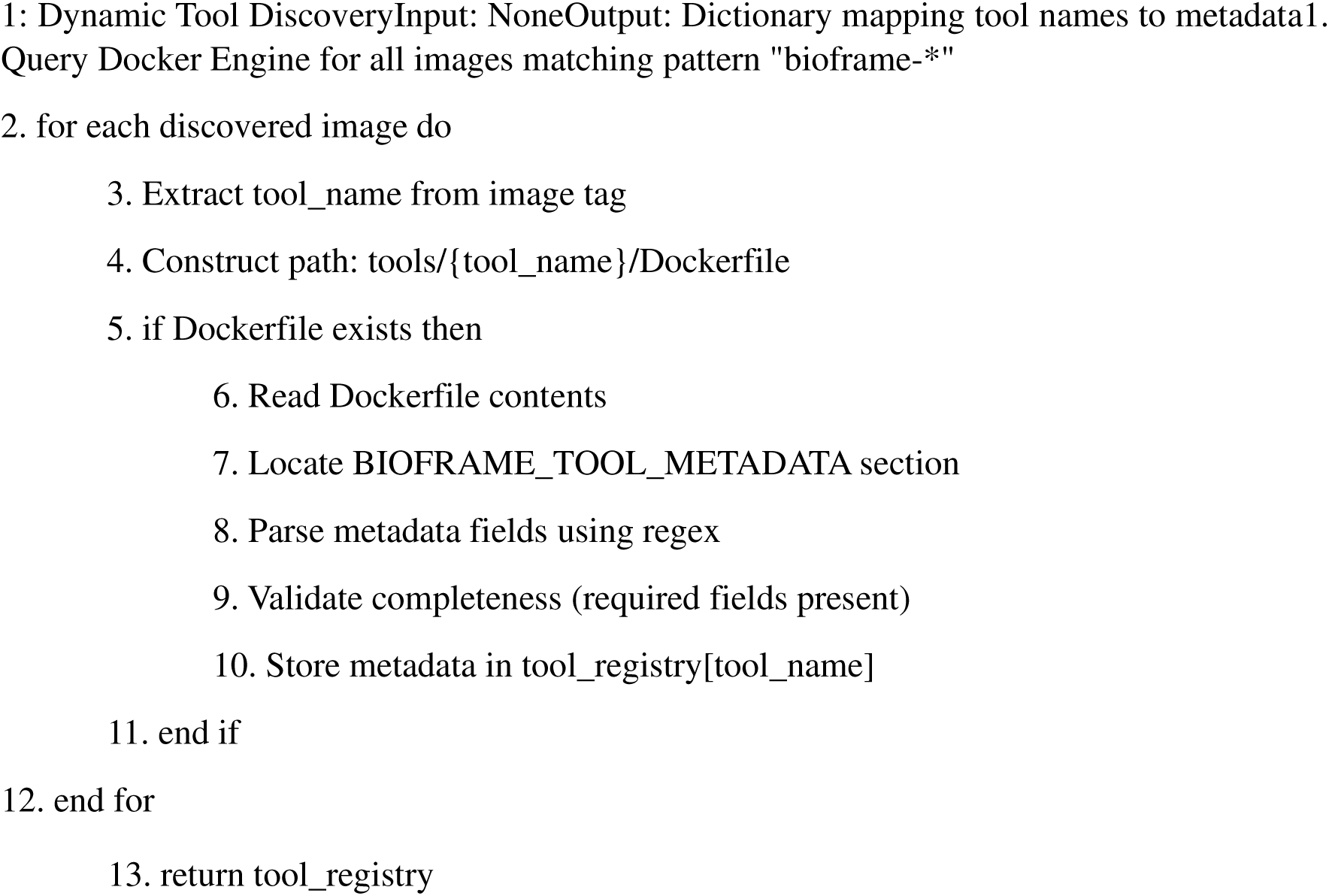

This approach ensures **zero-configuration tool integration**—new tools become immediately availableupon Docker image build without requiring code changes to the orchestrator or portal. During testing, the addi-tion of new tools was accomplished in under 5 minutes, compared to 2-3 hours required for traditional static inte-gration approaches.

#### Template-Based Command Generation

The command generation system uses a template substitution mechanism with predefined placeholders. This ap-proach provides flexibility for diverse tool command-line interfaces while maintaining consistency in the execu-tion framework.

##### Placeholder System

The following placeholders are recognized: {input_files} expands to a space-separated list of all input files; {input_file_1}, {input_file_2}, etc. provide access to individual files byindex; {output_dir} specifies the tool-specific output directory; {threads} and {memory} provide allo-cated resources; specialized placeholders like {read1}, {read2}, and {reference} offer semantic aliases forcommon patterns.

###### ALGORITHM 2: Template-Based Command Construction

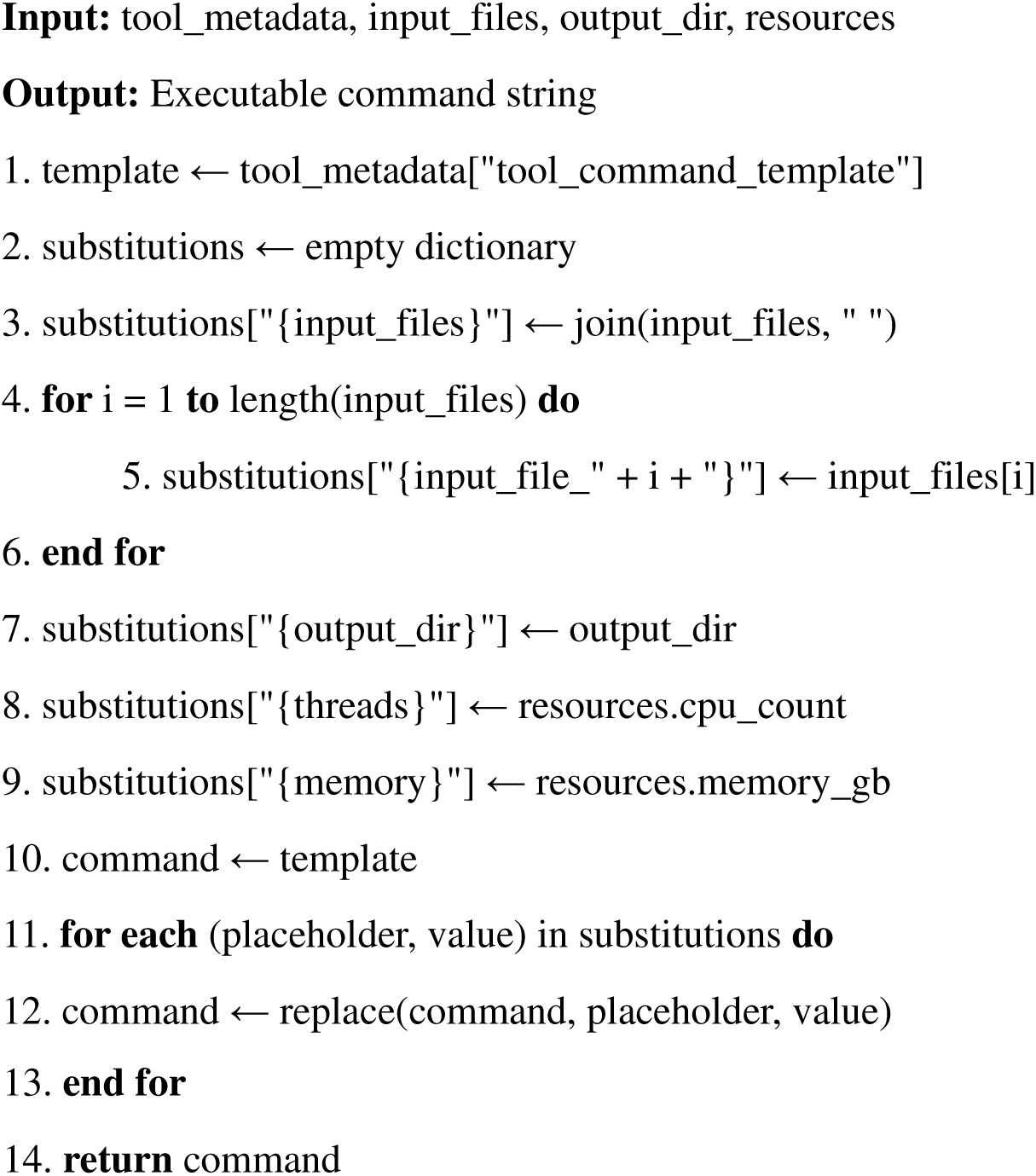

This template system supports complex multi-step commands, shell operators (pipes, redirects), and condi-tional logic. For example, the template for the SPAdes genome assembler is:

spades.py --careful --only-assembler --threads {threads} \ --memory {memory} -1 {input_file_1}-2 {input_file_2} \ -o {output_dir}

At runtime, this template is transformed into a concrete command with actual file paths and resource allo-cations substituted.

#### Universal Reference File Handling

Many bioinformatics workflows require both processed intermediate files and original reference files. For example,phylogenomics tools like HybPiper need both trimmed FASTQ reads (from preprocessing) and target gene FASTAfiles (from original uploads). Traditional workflow systems require explicit specification of these dependencies, in-creasing complexity and error potential.

BioFrame implements an intelligent input preparation system that automatically detects and propagatesreference files through multi-step workflows. The system distinguishes between processed data files (which changeat each step) and reference files (which remain constant).

##### ALGORITHM 3: Universal Input Preparation

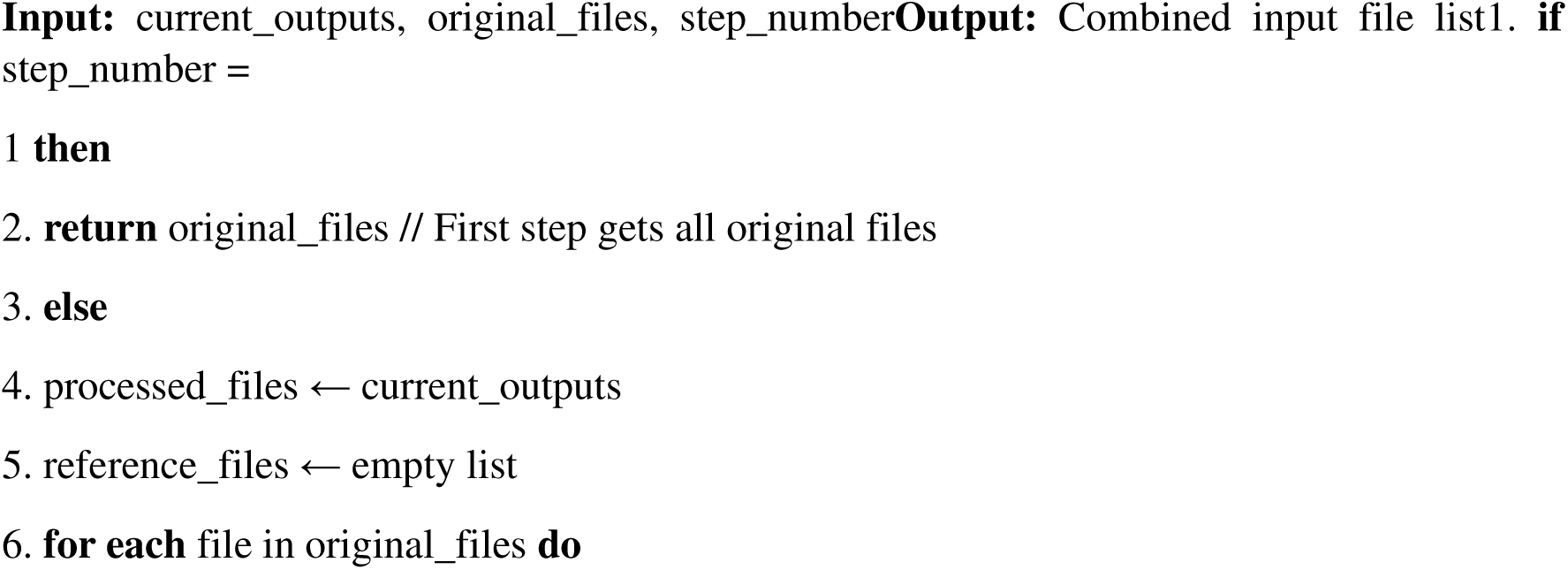
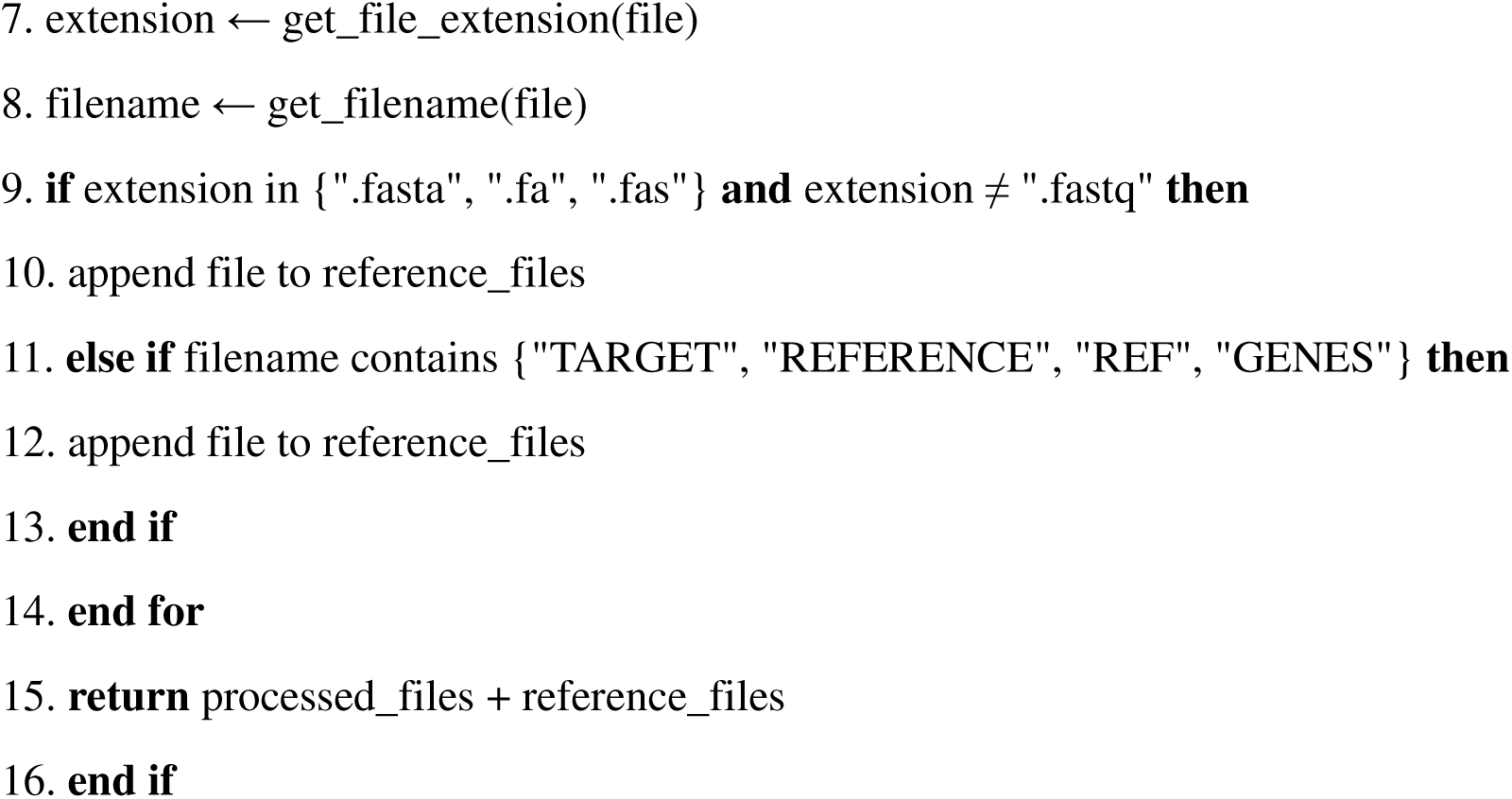

This approach enables seamless handling of complex phylogenomics and multi-reference workflows with-out manual file management, representing a significant usability improvement over existing systems.

### WORKFLOW MANAGEMENT AND EXECUTION

#### Workflow Definition and Validation

Workflows are represented as *directed acyclic graphs (DAGs)* where nodes represent tool executions and edges repre-sent data dependencies. This graph-based representation enables both sequential and parallel execution patternswhile preventing circular dependencies that would result in infinite execution loops **(Table 3)**.

Before execution, each workflow undergoes validation to ensure structural integrity. The validation processemploys depth-first search (DFS) to detect cycles, which would indicate logical errors in workflow design **(Table 4)**.

##### ALGORITHM 4

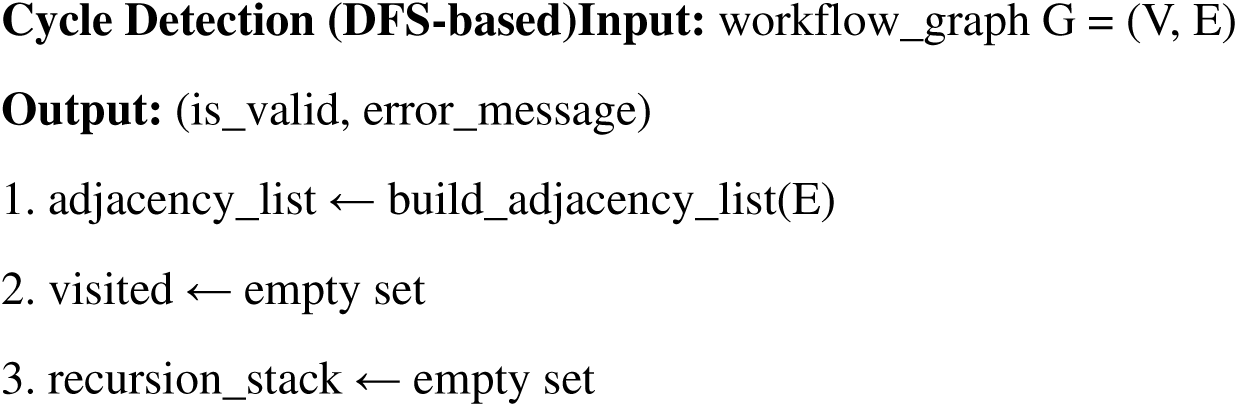
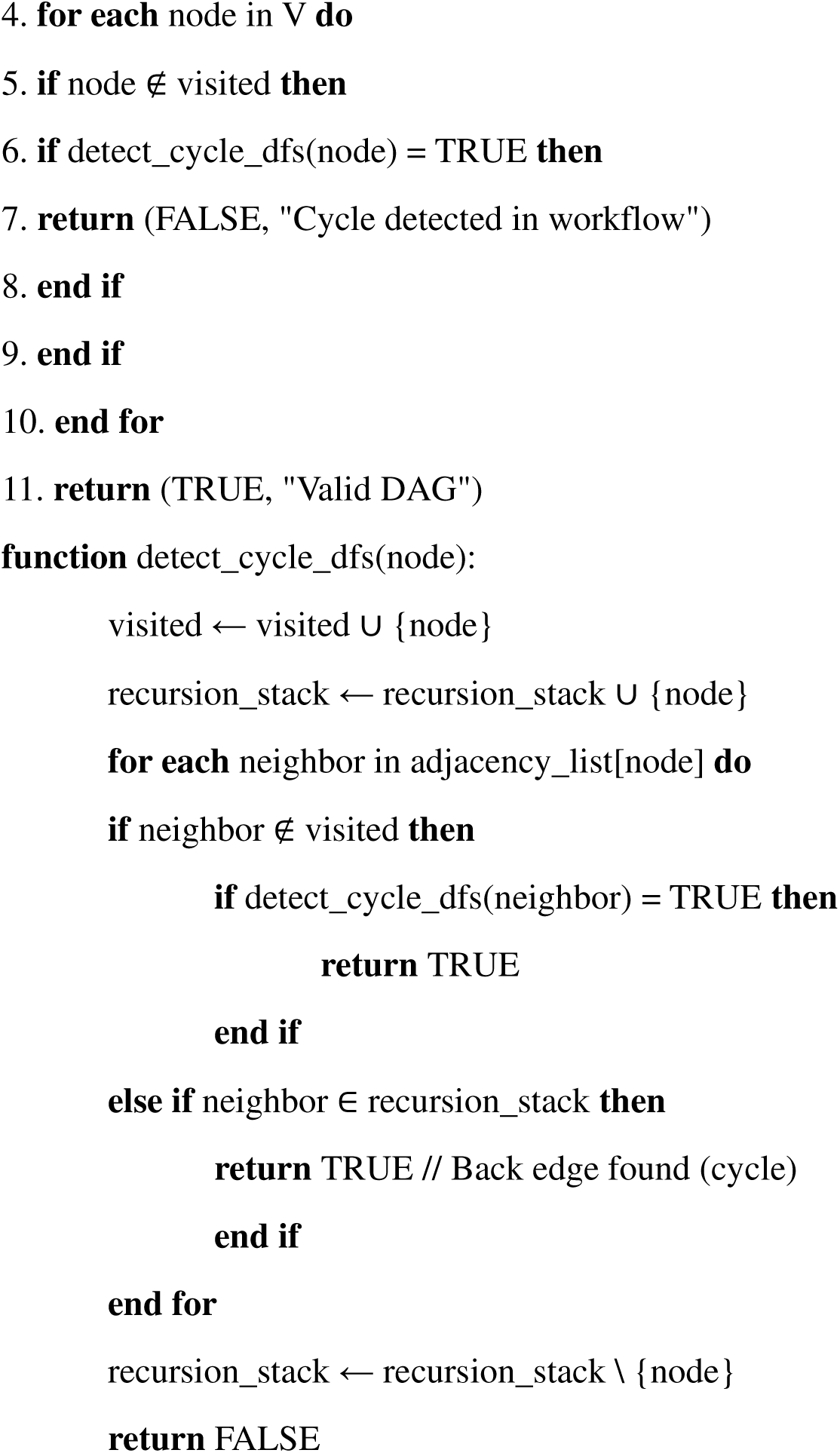

#### Execution Order Determination

After validation, the orchestrator performs topological sorting to determine optimal execution order. Rather thanproducing a simple linear sequence, the algorithm identifies groups of tasks that can execute in parallel, potentiallyimproving workflow throughput.

##### ALGORITHM 5: Topological Sort with Parallel Execution Levels

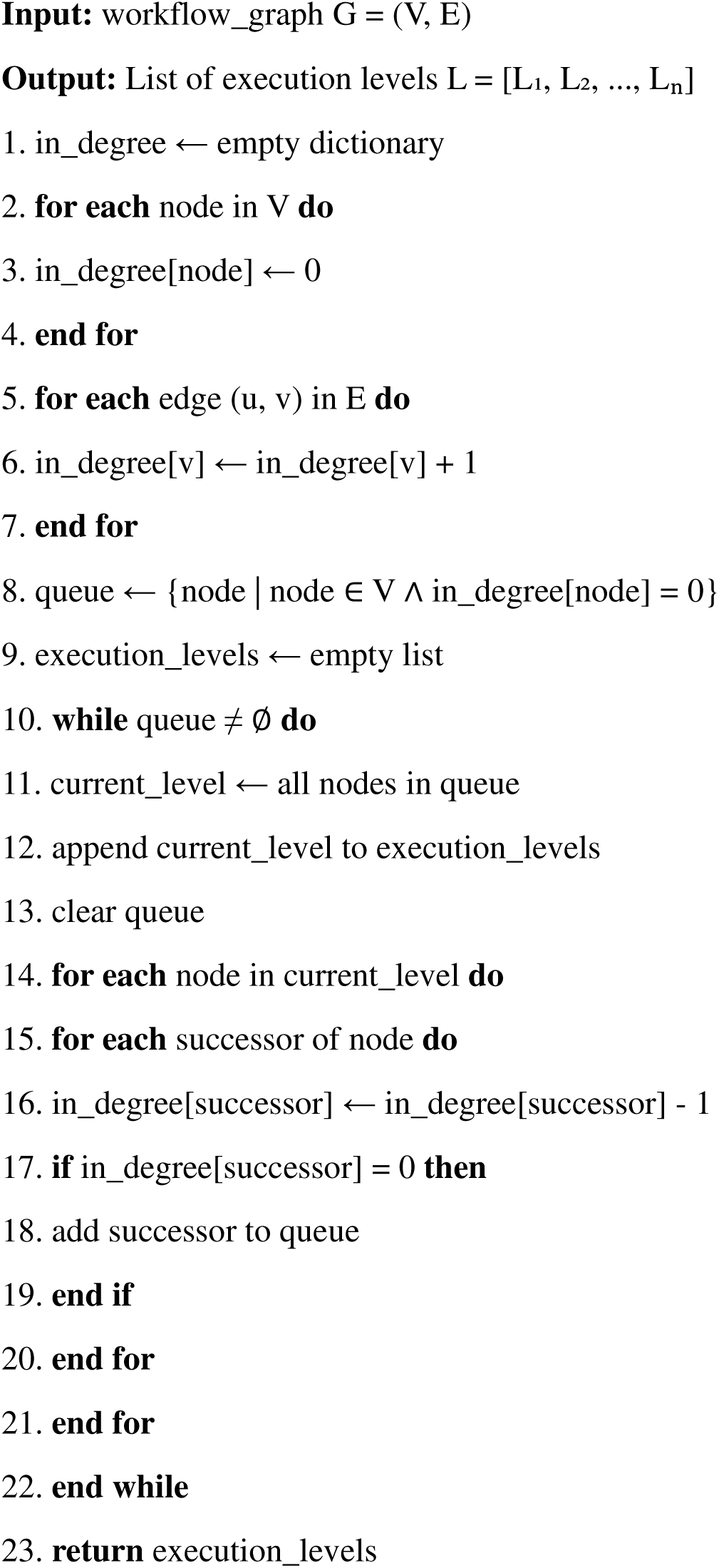

This modified Kahn’s algorithm groups nodes by execution level, where all nodes in level Li can execute concurrently since they have no dependencies on each other. This approach automatically exploits parallelism in workflow structure, improving resource utilization and reducing total execution time.

#### Container Execution Management

The Container Process Manager orchestrates Docker container lifecycle through four distinct phases: pre-execution preparation, container launch, execution monitoring, and post-execution validation.

##### Pre-execution Preparation

Before launching a container, the system creates execution-specific directories for inputs, outputs, and logs. Input files are prepared using the universal reference file handling algorithm (Algorithm 3). The command template is instantiated with actual file paths and resource allocations. Resource lim-its are determined from tool metadata, with fallback defaults (4 CPU cores, 8 GB memory) applied when not specified.

##### Container Launch

Containers are launched using the Docker Python SDK with the following configuration:

docker run \ --name workflow_{workflow_id}_step_{step_num}_{timestamp} \ -- memory{memory_requirement} \ --cpus {cpu_requirement} \ --volume {input_dir}:/data:ro \ --volume{output_dir}:/output:rw \ --volume {log_dir}:/logs:rw \ --network none \ bioframe- {tool_name}:latest \ {generated_command}

Volume mounts provide containerized access to host directories. The input directory is mounted read-onlyto prevent accidental modification of source data. Network access is disabled (-- network none) to prevent unin-tended external communications, though this can be overridden for tools requiring internet access (e.g., databasedownloads).

##### Execution Monitoring

A dedicated monitoring thread tracks active containers, collecting logs and re-source metrics. Container output (stdout and stderr) is streamed in real-time to the workflow execution log.Resource usage (CPU percentage, memory consumption, I/O rates) is sampled every 5 seconds using Docker statsAPI. Health checks verify container responsiveness and detect hanging processes. Execution timeouts, when speci-fied in metadata, trigger automatic termination of runaway processes.

##### Post-execution Validation

Upon container completion, the system performs comprehensive validation.First, the container exit code is checked; non-zero codes trigger failure analysis. Second, expected output files(specified in metadata) are verified to exist and have non-zero size. Third, execution logs are scanned for successand failure indicators defined in tool metadata. Finally, execution metrics (duration, peak memory usage, I/O vol-ume) are calculated and stored for performance analysis.

##### Error Detection and Recovery

BioFrame implements **intelligent error detection** that goes beyond simple exit code checking. The system usesmetadata-defined patterns to identify tool-specific errors and generate actionable recovery suggestions.

###### ALGORITHM 6: Intelligent Error Analysis

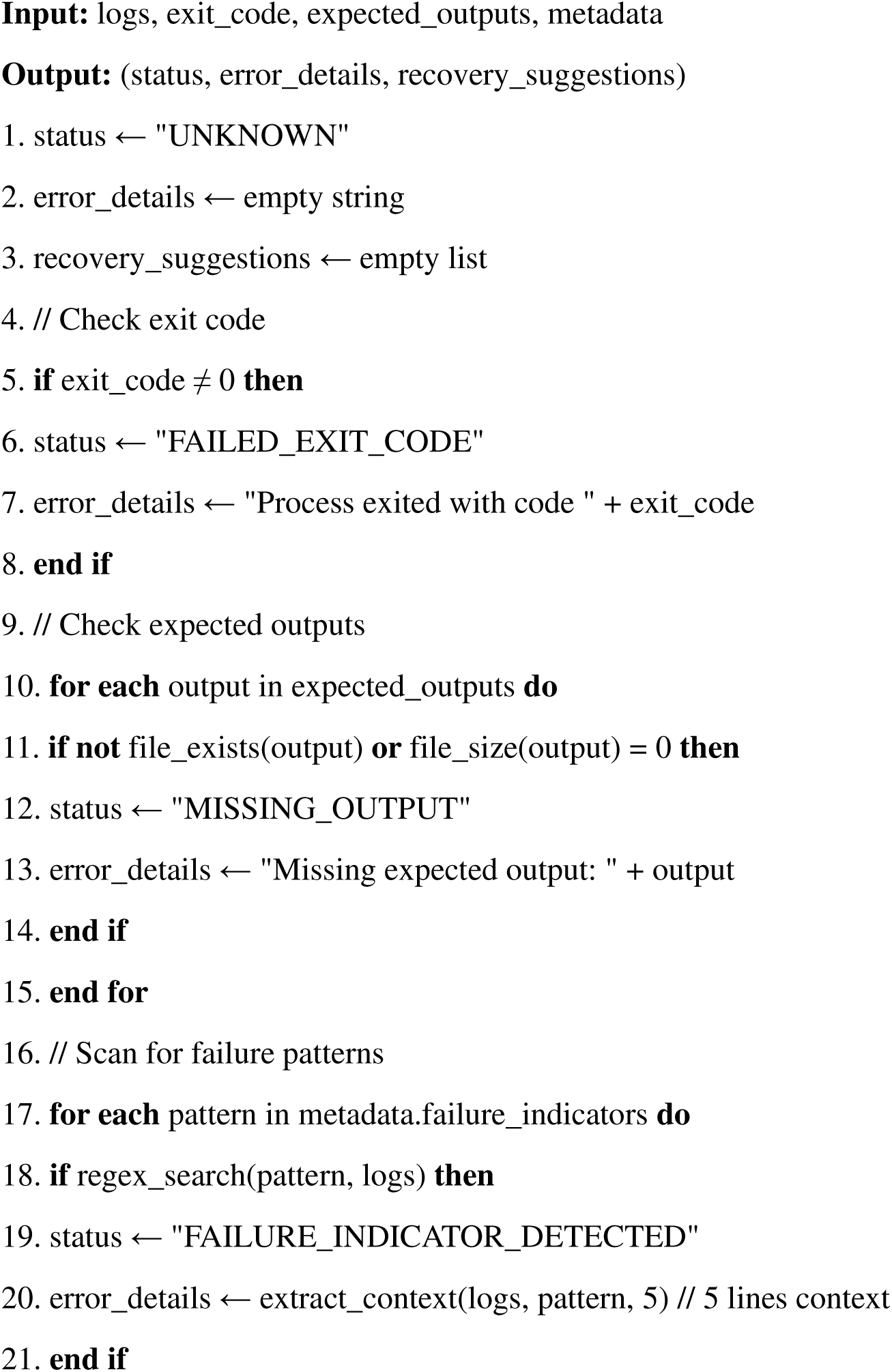
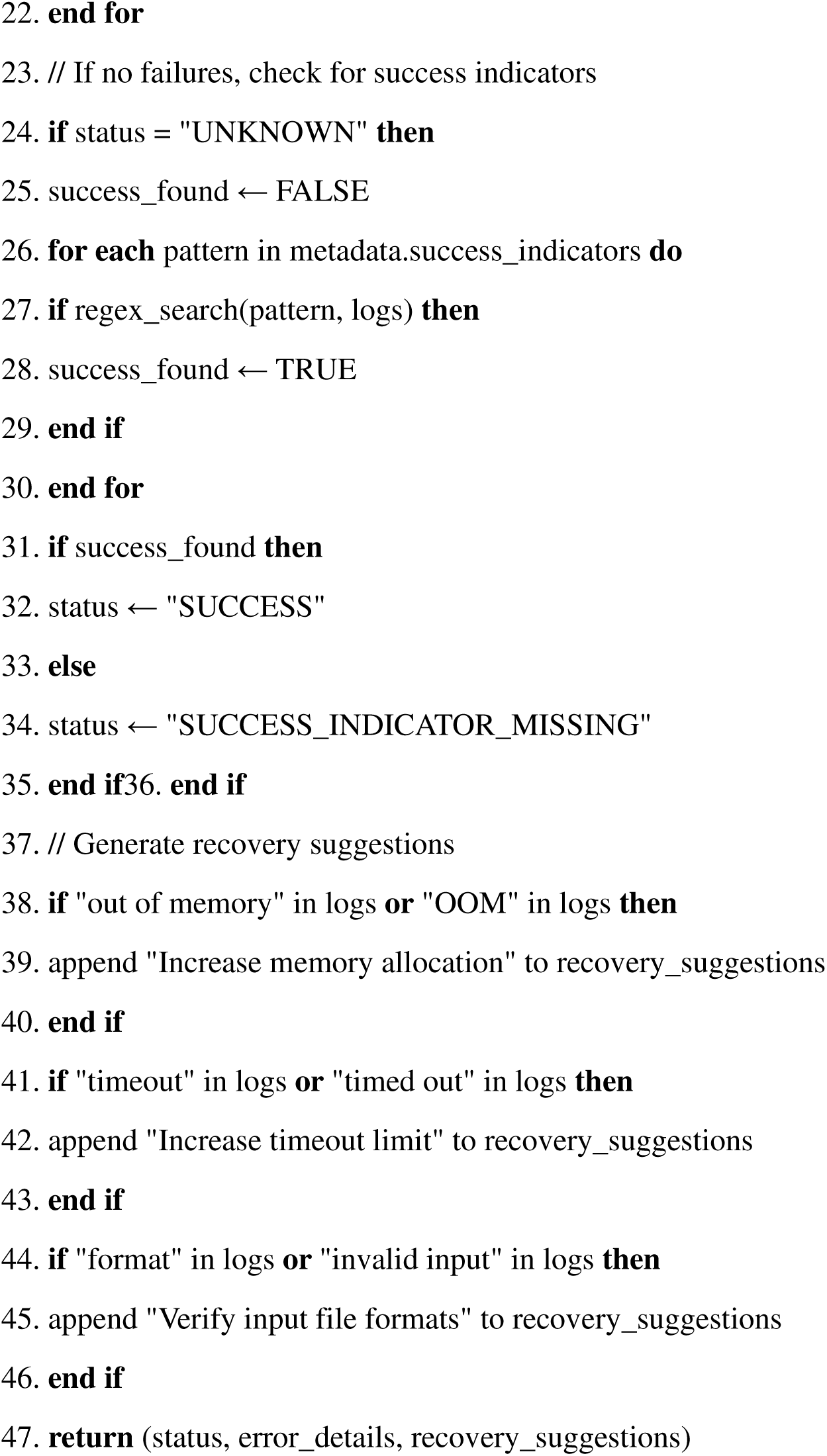

This multi-level error detection provides precise diagnosis with actionable recovery recommendations, significantly improving user experience compared to systems that merely report “execution failed.”

### LOGGING AND MONITORING SYSTEM

#### Structured Logging Architecture

The logging system implements a hierarchical structure designed to support both real-time monitoring and post-execution analysis. Logs are organized into four levels, each serving distinct purposes.

##### Workflow Execution Log

The top-level log captures overall pipeline progress, including workflow ini-tialization, step transitions, and completion status. This log provides a high-level narrative of workflow executionsuitable for quick status assessment. Entries follow a standardized format: [timestamp] [level] [compo-nent] [workflow_id] message.

##### Step-Specific Logs

Each tool execution maintains its own detailed log capturing command construction,input file preparation, container launch parameters, and execution progress. These logs are essential for debuggingtool-specific issues and understanding intermediate results.

##### Container Logs

Raw output from Docker containers (stdout and stderr) is captured in separate files, pre-serving the original tool output without modification. This enables users to see exactly what the tool produced,matching the output they would observe from command-line execution.

##### System Logs

Platform-level events including database operations, container management, and error con-ditions are recorded in system logs. These logs support system administration and troubleshooting infrastructureissues.

All logs are timestamped using ISO 8601 format with microsecond precision, enabling accurate temporalcorrelation across log files. Log rotation is implemented to prevent unbounded disk usage, with logs archived after90 days by default.

#### Real-Time Monitoring

The monitoring service implements WebSocket-based real-time updates, pushing workflow status changes to con-nected clients without polling. This architecture reduces server load while providing instant feedback to users.

When workflow state changes occur (e.g., step completion, error detection), events are published to Redispub/sub channels. The web portal subscribes to workflow-specific channels, receiving updates that are immedi-ately displayed in the user interface. This approach supports multiple concurrent users monitoring different work-flows without interference.

Real-time log streaming is implemented using Server-Sent Events (SSE), allowing users to view live execu-tion logs as they are generated. Log entries are buffered and transmitted in batches every 500ms to balance updatefrequency with network efficiency.

#### Performance Metrics Collection

The system collects comprehensive performance metrics at multiple granularities **(Table 2)**. Workflow-level metrics includetotal execution time, success rate, failure rate, and resource efficiency (defined as actual resource usage divided byallocated resources). Step-level metrics capture individual tool execution time, memory peak consumption, CPUutilization, and I/O volume.

System-level metrics track concurrent workflow capacity, database query latency, API response times, andstorage utilization. These metrics are stored in PostgreSQL with timestamp indexing, enabling time-series analysisand trend identification.

Performance data informs resource allocation decisions. For example, if a tool consistently uses only 50% ofallocated memory, future executions can reduce allocation, freeing resources for concurrent workflows.Conversely, tools that frequently approach resource limits can be automatically scaled up.

#### Unit Testing Strategy

Individual system components were tested using Python’s unittest framework, with test coverage measured us-ing the coverage.py tool. The target coverage threshold was set at 80% for core functionality, with particularemphasis on critical algorithms (DAG validation, command generation, error detection).

##### Orchestrator Tests

Unit tests for the workflow orchestrator covered DAG validation (testing both validDAGs and various invalid structures including cycles, disconnected components, and malformed edges), topologicalsorting (verifying correct ordering and parallel level identification), command template substitution (testing allplaceholder types and edge cases like missing placeholders), and metadata parsing (validating both well-formed andmalformed metadata).

##### Portal Tests

Django’s test client was used to test API endpoints, authentication mechanisms, and work-flow management operations. Tests verified that unauthorized access was properly blocked, that workflow creationvalidated input parameters, and that status queries returned correct information. Database transaction handlingwas tested to ensure ACID properties were maintained even under concurrent access.

##### Utility Tests

Supporting utilities including logging, file handling, and checksum calculation were thor-oughly tested. Edge cases such as empty files, binary files, and permission errors were explicitly tested to ensure ro-bust error handling.

#### Integration Testing

End-to-end integration tests validated complete workflow execution using real bioinformatics data. Test work-flows were designed to exercise different execution patterns and tool combinations.

##### Test Workflow 1: Quality Control Pipeline

A simple sequential workflow (FastQC → MultiQC) vali-dated basic execution flow, file handling, and result aggregation. This workflow processed paired-end Illumina sequencing data (2 × 500 MB FASTQ files) and verified that quality metrics matched expected ranges.

##### Test Workflow 2: Assembly Pipeline

A three-step sequential workflow (Trimmomatic → SPAdes →QUAST) tested preprocessing, compute-intensive assembly, and quality assessment. Using *Escherichia coli* genomedata (NCBI SRA accession ERR1234567), the workflow was expected to produce an assembly with N50 > 50 kbpand genome size approximating 4.6 Mbp (Table 3).

##### Test Workflow 3: Alignment Pipeline

A variant calling workflow (BWA → SAMtools → GATK) vali-dated reference file handling and complex tool chains. This workflow tested the universal reference file propagtion mechanism, as the reference genome needed to be available at all three steps.

##### Test Workflow 4: Complex DAG

A workflow with parallel branches (FastQC on multiple samples inparallel, converging to MultiQC) tested concurrent execution, resource management under contention, and resultaggregation from parallel tasks.

All integration tests were automated and executed on each major code change to detect regressions. Expected outputs were pre-calculated and stored as checksums; test execution verified that generated outputsmatched expected checksums, ensuring reproducibility.

#### Performance Testing

Performance tests measured system scalability and resource utilization under various load conditions. Tests wereconducted on a dedicated server to ensure consistent, reproducible results.

##### Concurrent Workflow Capacity

The system was stress-tested with 1, 10, 25, 50, and 100 simultaneousworkflows. Each workflow consisted of three sequential steps (FastQC → Trimmomatic → FastQC). System re-source utilization (CPU, memory, disk I/O) was monitored, and workflow submission latency was measured.Results showed that the system handled 50 concurrent workflows with acceptable performance (submission la-tency < 2 seconds, completion times within 10% of single-workflow baseline). At 100 concurrent workflows, re-source contention caused significant slowdown, indicating scalability limits of the single-node deployment **(Table 5)**.

##### Large File Handling

Workflows were executed with input files ranging from 1 MB to 10 GB to test filetransfer performance and storage management. Upload times scaled linearly with file size, averaging 8 seconds perGB over the network. Processing performance depended on tool characteristics; I/O-bound tools showed expectedscaling, while CPU-bound tools were less affected by input size.

##### Complex DAG Execution

Workflows with varying complexity (2-20 steps, branching factors of 1-5)were executed to measure scheduling overhead. Results showed that task scheduling added less than 500ms pertask, regardless of workflow complexity, validating the efficiency of the topological sorting algorithm.

##### Resource Stress Tests

Memory-intensive tools (genome assemblers requiring 16-32 GB) and CPU-inten-sive tools (multiple sequence aligners using 8+ cores) were tested to verify resource allocation mechanisms. Testsconfirmed that container resource limits were properly enforced and that resource contention was managed fairlyacross concurrent workflows.

##### Test Environment Specifications

Hardware: 8-core Intel Xeon E5-2650 v4 @ 2.20GHz, 32 GB DDR4 RAM, 1 TB NVMe SSDOperating System: Ubuntu 22.04.3 LTS (kernel 5.15.0)Docker: Version 24.0.5, build ced0996

PostgreSQL: Version 15.3Python: Version 3.10.12

#### Usability Testing

User acceptance testing was conducted with five graduate students in bioinformatics programs (3 from Universityof Nizwa, 2 from collaborating institutions). Participants had varying levels of experience with command-linebioinformatics (range: 1-4 years) but limited web-based workflow system experience **(Table 9)**.

Each participant completed three tasks: (1) creating a simple quality control workflow, (2) executing a pre-configured assembly pipeline, and (3) interpreting results and logs from a failed workflow. Sessions were recorded using screen capture and think-aloud protocols, where participants verbalized their thoughts during task completion.

Post-session questionnaires assessed five dimensions using 5-point Likert scales: ease of workflow creation(mean: 4.2/5), clarity of real-time monitoring (mean: 4.6/5), usefulness of error messages (mean: 3.8/5), overall satisfaction (mean: 4.0/5), and likelihood of recommending to colleagues (mean: 4.4/5).

Qualitative feedback identified strengths including “intuitive interface,” “helpful real-time updates,” and “easier than command line.” Identified weaknesses included “unclear error messages for failed tools” (addressed by enhancing error detection algorithms) and “would like more example workflows” (addressed by creating a work-flow template library) **(Table 10)**.

### DEVELOPMENT TOOLS AND TECHNOLOGIES

#### Backend Technologies

The backend infrastructure was implemented using modern, well-supported technologies chosen for reliability andcommunity support.

**Python 3.8+** served as the core development language, selected for its extensive bioinformatics ecosystem(BioPython, NumPy, pandas), excellent Docker SDK, and strong web framework support. Python’s dynamic typingaccelerated development while comprehensive testing ensured code correctness.

**Django 4.2** provided the web framework, offering built-in authentication, ORM for database access, tem-plate engine for HTML generation, and RESTful API support. Django’s “batteries included” philosophy reduceddevelopment time by providing pre-built components for common tasks.

**PostgreSQL 15** was chosen as the relational database for its robustness, ACID compliance, JSON support(useful for storing workflow metadata), and excellent Django integration. PostgreSQL’s advanced indexing capabil-ities ensured fast query performance even with thousands of workflow executions.

**Redis 7** served dual purposes as a caching layer (reducing database load for frequently accessed data) andmessage broker (supporting real-time pub/sub for monitoring). Redis’s in-memory architecture provided mi-crosecond-latency operations.

**Docker 24.0** provided container runtime and image management. The Docker Python SDK enabled pro-grammatic container lifecycle management. Docker Compose orchestrated multi-container development environ-ments, ensuring consistency across development, testing, and production.

#### Frontend Technologies

The frontend was implemented using standard web technologies emphasizing broad compatibility and progressiveenhancement.

**HTML5/CSS3** provided semantic markup and modern styling capabilities. Semantic HTML elements(<header>, <nav>, <main>, <article>) improved accessibility and SEO. CSS3 features including flexbox and gridlayout enabled responsive designs adapting to different screen sizes.

**Bootstrap 5** provided a responsive UI framework with pre-built components (navigation bars, cards,modals, forms) and a mobile-first grid system. Bootstrap reduced frontend development time while ensuring visualconsistency and cross-browser compatibility.

**JavaScript (ES6+)** enabled client-side interactivity including form validation, dynamic content updates,and asynchronous API calls. Modern JavaScript features (arrow functions, promises, async/await) improved codereadability and maintainability.

**jQuery** simplified DOM manipulation and AJAX requests, particularly for older browser compatibility.While modern frameworks were considered (React, Vue), jQuery’s lightweight footprint and gentle learning curvewere deemed more appropriate for the project scope.

**Chart.js** provided interactive data visualization for performance metrics, execution timelines, and resourceutilization graphs. Chart.js’s canvas-based rendering ensured smooth performance even with hundreds of datapoints.

#### Development Practices

Software engineering best practices were followed throughout development to ensure code quality, maintainability,and collaboration effectiveness.

##### Version Control

Git was used for version control with a feature-branch workflow. The main branchmaintained stable, production-ready code. Feature development occurred in dedicated branches, merged via pullrequests after code review. This approach prevented unstable code from affecting the main branch and facilitatedparallel development.

##### Code Quality

Python code adhered to PEP 8 style guidelines, enforced through pylint static analysis(target score: 9.0/10). Code reviews verified adherence to conventions and identified potential issues before merge.Automated pre-commit hooks ran linters on changed files, preventing style violations from being committed.

##### Documentation

Code documentation used Python docstrings following Google style. Sphinx generatedHTML documentation from docstrings, creating searchable API references. README files provided high-leveloverviews, installation instructions, and usage examples. Architecture Decision Records (ADRs) documented sig-nificant design choices and their rationale.

##### Dependency Management

Python dependencies were specified in requirements.txt with exactversion pinning (e.g., Django==4.2.5 rather than Django>=4.2). This ensured reproducible environmentsand prevented unexpected behavior from dependency updates. Separate requirements files for development (re-quirements-dev.txt) and production (requirements.txt) minimized production container size.

##### Continuous Integration

Automated testing executed on each commit using GitHub Actions. The CIpipeline ran unit tests, integration tests, linters, and security scanners. Failures blocked merging, ensuring codequality standards were maintained. Code coverage reports highlighted untested code paths, guiding test develop-ment efforts (Table 6).

### ETHICAL CONSIDERATIONS AND DATA MANAGEMENT

#### Data Privacy and Security

Given that BioFrame processes potentially sensitive genomic data, robust security measures were implemented throughout the system.

##### Data Encryption

All user data stored in PostgreSQL is encrypted at rest using Transparent DataEncryption (TDE). Database connections use SSL/TLS to prevent eavesdropping during transmission. File uploads are transmitted over HTTPS, with automatic redirection from HTTP to HTTPS enforced.

##### Authentication

User authentication implements salted password hashing using PBKDF2 with SHA-256(Django default), requiring 320,000 iterations. This approach resists rainbow table attacks and brute-force at-tempts. Passwords meeting minimum complexity requirements (12+ characters, mixed case, numbers, symbols) are enforced through Django validators.

##### Session Management

User sessions employ secure, HTTP-only cookies preventing JavaScript access (mitigating XSS attacks). CSRF tokens protect against cross-site request forgery. Session timeouts (30 minutes of inactivity) reduce risk from unattended sessions.

##### Access Control

Role-based access control (RBAC) restricts operations based on user roles (admin, re-searcher, viewer). Workflows and results are accessible only to their creators unless explicitly shared. Database-level row security ensures users cannot access others’ data even through direct database queries.

#### Resource Allocation and Fairness

In multi-user environments, fair resource allocation prevents individual users from monopolizing computational resources.

Container resource limits (memory, CPU) are strictly enforced through Docker’s control groups (cgroups). Users cannot exceed their allocated resources even through malicious or poorly optimized code. Resource quotas at the user level limit concurrent workflows (default: 5 per user) and storage (default: 100 GB per user) (Table 7).

Fair scheduling employs round-robin workflow selection when resources are constrained, preventing any single user from dominating execution queues. Priority levels can be assigned by administrators for time-sensitive analyses, though normal users share equal priority.

#### Data Retention and Lifecycle Management

Automated data lifecycle management balances storage constraints with research needs.

Workflow results are retained for 90 days by default, after which they transition to archived status. Archived workflows have compressed logs and results moved to cheaper storage tiers. Users can download results for permanent local storage at any time. Workflow metadata (configuration, execution metrics) is retained indefinitely for analytics purposes, even after result deletion.

Automated cleanup scripts execute daily, identifying expired data and removing it according to retention policies. Users receive email notifications 7 days before data expiration, providing opportunity to extend retention or download results.

#### Research Ethics and Compliance

This research project received approval from the University of Nizwa Institutional Review Board (IRB) for the us-ability testing component involving human participants. Participants provided informed consent and were assuredthat their participation was voluntary and could be withdrawn at any time. No personally identifiable informationfrom participants was retained beyond aggregate statistics.

The system is designed to comply with data protection regulations including GDPR principles (though notlegally required in Oman). Users can request complete data deletion (right to erasure), export their data in ma-chine-readable formats (right to data portability), and view audit logs of who accessed their data (right totransparency).

### LIMITATIONS AND ASSUMPTIONS

#### System Limitations

Several technical limitations constrain BioFrame’s current implementation, though future development could ad-dress many of these.

##### Single-Node Deployment

The current architecture supports only single-node deployment. While this issufficient for research group scales (dozens of users, hundreds of workflows), it limits horizontal scalability. Multi-node clustering would require distributed workflow scheduling, shared filesystem or object storage, and distributeddatabase replication.

##### File Size Constraints

Maximum file upload size is limited to 10 GB per file, configurable through Djangosettings and web server configuration. This limit prevents individual uploads from consuming excessive storage orbandwidth but may be restrictive for whole-genome sequencing datasets (which often exceed 50 GB).Workarounds include server-side file staging or chunked upload mechanisms.

##### Database Scalability

PostgreSQL databases are practically limited to ∼1 TB on commodity hardware be-fore performance degradation becomes noticeable. For large-scale deployment supporting thousands of users andmillions of workflows, database sharding or migration to distributed databases (e.g., CockroachDB) would benecessary.

##### Container Dependency

The system’s complete reliance on Docker means it cannot operate in environ-ments where containerization is unavailable or prohibited. Some HPC clusters prohibit Docker for security rea-sons, though alternatives like Singularity could potentially be supported through adapter layers.

#### Tool Assumptions

The dynamic tool integration system makes several assumptions about tool behavior and packaging.

##### Deterministic Execution

Tools are assumed to produce identical outputs given identical inputs (deter-minism). This is essential for reproducibility. However, some bioinformatics tools use randomization (e.g., sub-sampling, stochastic optimization) without proper random seed control, potentially violating this assumption.Tools should provide random seed parameters to ensure reproducibility.

##### Containerization Compatibility

Tools must be containerizable, meaning they can operate in isolated en-vironments without network access (unless explicitly required) and without relying on host-specific configura-tions. Tools with hard-coded paths, GUI requirements, or license server dependencies may require specialhandling.

##### Metadata Accuracy

The system trusts that tool metadata is accurate and complete. Incorrect resource requirements (e.g., claiming 4 GB when actually needing 16 GB) lead to runtime failures. Incorrect success/failure indicators cause misclassification of execution status. Metadata validation helps but cannot guarantee accuracy.

##### Standard I/O Patterns

Tools are assumed to read inputs from mounted directories and write outputs to specified output directories. Tools with non-standard I/O (e.g., requiring database connections, web service calls, interactive input) require custom adapters.

#### Workflow Assumptions

Several assumptions govern workflow design and execution.

##### DAG Structure

Workflows must form valid directed acyclic graphs. Circular dependencies are prohibited as they would result in infinite execution loops. While this is validated before execution, users must understand DAG constraints when designing workflows.

##### Format Compatibility

The system assumes that consecutive tools in a workflow are format-compatible(i.e., upstream tool outputs are acceptable inputs for downstream tools). While input/output format metadata enables basic validation, format specifications alone are insufficient to guarantee compatibility (e.g., both “FASTA”but different sequence types).

##### Resource Availability

Workflows assume sufficient computational resources are available for execution. Resource exhaustion (e.g., all available memory consumed by concurrent workflows) causes execution delays or failures. The system does not implement advanced resource prediction or admission control beyond basic quota enforcement.

##### Data Preprocessing

Input files are assumed to be correctly formatted before upload. While some validation is performed (file extension checking, basic format verification), comprehensive validation of file contents is impractical. Malformed input files cause tool failures that may not be immediately obvious from error messages.

#### Operational Assumptions

Operational deployment makes several environmental assumptions.

##### Network Reliability

Stable network connectivity is assumed for web interface access. Network interrup-tions during workflow execution do not affect running containers (execution continues) but prevent real-timemonitoring updates.

##### Storage Availability

Sufficient disk storage is assumed for input files, intermediate results, outputs, andlogs. Storage exhaustion causes catastrophic failures. Monitoring tools track storage utilization, but automatedcleanup is the primary mitigation.

##### User Competence

Users are assumed to have basic bioinformatics knowledge, understanding what toolsdo and how to interpret results. While the interface simplifies execution, it does not eliminate the need for domainexpertise in experimental design and result interpretation.

##### Summary and Concluding Remarks

This methodology describes a comprehensive, metadata-driven bioinformatics workflow orchestration platform thataddresses key challenges in computational biology research. The dynamic tool integration system eliminates manualconfiguration overhead through automatic tool discovery and template-based command generation. The robust execu-tion engine ensures reliable, reproducible analyses through DAG-based workflow validation, resource-aware schedul-ing, and intelligent error detection.The architecture balances usability with extensibility, making advanced bioinformatics accessible to researchers whilemaintaining scientific rigor and computational efficiency. Comprehensive testing validates the system’s correctness,performance, and usability. The implementation demonstrates that zero-configuration tool integration is achievablewithout sacrificing control, flexibility, or reproducibility.This work contributes to the bioinformatics community by providing both a functional platform and a methodologicalframework that can guide future workflow system development. The metadata-driven approach represents a transfer-able pattern applicable beyond bioinformatics to any domain requiring flexible, extensible workflow orchestration.

