## Supplementary material for "BioFrame: Enhancing Reproducibility and Accessibility in Genomics through Web-Based Workflow Design": Tables

### BioFrame Data Tables

Comprehensive Data for Academic Essay Submission

University of Nizwa | Department of Biological Science | 2024-2025

### Table 1: Integrated Bioinformatics Tools in BioFrame Platform

*Complete catalog of containerized bioinformatics tools integrated into the BioFrame platform, organized by functional category with version information and primary applications.*

| Category | Tool Name | Version | Container Image | Primary Application |
|---|---|---|---|---|
| Quality Control | FastQC | 0.11.9 | bioframe-fastqc:latest | Sequencing data quality assessment |
| Quality Control | MultiQC | 1.14 | bioframe-multiqc:latest | Aggregate quality reports |
| Read Processing | Trimmomatic | 0.39 | bioframe-trimmomatic:latest | Read trimming and filtering |
| Read Processing | Seqtk | 1.3 | bioframe-seqtk:latest | Sequence processing toolkit |
| Genome Assembly | SPAdes | 3.15.5 | bioframe-spades:latest | De novo genome assembly |
| Assembly QC | QUAST | 5.2.0 | bioframe-quast:latest | Assembly quality assessment |
| Assembly Improvement | Pilon | 1.24 | bioframe-pilon:latest | Assembly polishing and correction |
| Read Alignment | BWA | 0.7.17 | bioframe-bwa:latest | Burrows-Wheeler alignment |
| Alignment Processing | SAMtools | 1.17 | bioframe-samtools:latest | SAM/BAM file manipulation |
| Genome Analysis | BEDtools | 2.30.0 | bioframe-bedtools:latest | Genomic interval operations |
| Variant Calling | GATK | 4.2.0.0 | bioframe-gatk:latest | Variant discovery and genotyping |
| Phylogenomics | HybPiper | 2.1.6 | bioframe-hybpiper:latest | Target enrichment gene recovery |
| Sequence Alignment | MAFFT | 7.505 | bioframe-mafft:latest | Multiple sequence alignment |
| Alignment Trimming | trimAl | 1.4.1 | bioframe-trimal:latest | Alignment trimming |
| Phylogenetic Trees | IQ-TREE | 2.2.0 | bioframe-iqtree:latest | Maximum likelihood phylogenetics |
| Phylogenetic Trees | RAxML | 8.2.12 | bioframe-raxml:latest | ML phylogenetic inference |
| Phylogenetic Trees | FastTree | 2.1.11 | bioframe-fasttree:latest | Fast approximate ML trees |
| Species Trees | ASTRAL | 5.7.8 | bioframe-astral:latest | Species tree reconstruction |

*All tools containerized using Docker with Ubuntu 22.04 LTS base images. Version numbers represent the bioinformatics tool version, not container version.*

#### Table 2: System Performance Metrics

*Performance benchmarks measured on test hardware (8-core Intel Xeon E5-2650 v4, 32 GB RAM, 1 TB NVMe SSD, Ubuntu 22.04 LTS).*

| Metric Category | Metric | Measured Value | Significance |
|---|---|---|---|
| Workflow Management | Workflow submission latency | `< 2 seconds` | Time from user submission to execution start |
| Workflow Management | Workflow configuration parsing | `< 500 ms` | YAML workflow parsing and validation |
| Execution Scheduling | Task scheduling overhead | `< 500 ms per task` | Time to prepare and launch each pipeline step |
| Execution Scheduling | Container startup time | `2-5 seconds` | Docker container initialization time |
| Scalability | Concurrent workflow capacity | `50+ workflows` | Maximum simultaneous workflows on test hardware |
| Scalability | Optimal concurrent workflows | `10-15 workflows` | Concurrent workflows without performance degradation |
| Database Performance | Metadata query response | `< 100 ms` | Average PostgreSQL query time |
| Database Performance | Workflow status update | `< 50 ms` | Database write operation latency |
| Web Interface | API response time (avg) | `< 200 ms` | REST API endpoint response time |
| Web Interface | Page load time | `< 1.5 seconds` | Full page render time |
| File Management | File upload throughput | `~8 seconds/GB` | Network transfer rate for input files |
| File Management | Maximum file upload size | `10 GB per file` | Configurable upload size limit |
| Logging System | Log streaming latency | `< 1 second` | Real-time log update delay |
| Resource Monitoring | Resource monitoring interval | `5 seconds` | Container CPU/memory sampling frequency |
| Storage | Storage scalability | `Petabyte-scale` | Limited by storage backend capacity |

*Performance metrics represent average values across 100+ workflow executions. Individual workflow performance varies based on tool requirements and data volume.*

#### Table 3: E. coli Genome Assembly Pipeline - Execution Results

*Results from integration testing using Escherichia coli K-12 paired-end Illumina sequencing data (2 × 500 MB FASTQ files, ~100× coverage).*

| Pipeline Step | Tool | Execution Time | Peak Memory | CPU Utilization | Key Outputs |
|---|---|---|---|---|---|
| Step 1: Quality Control | FastQC | 3 min 24 sec | 2.1 GB | 85% | HTML reports, Q30 score: 92.3% |
| Step 2: Read Trimming | Trimmomatic | 8 min 16 sec | 3.8 GB | 78% | Trimmed FASTQ, 94.2% reads retained |
| Step 3: De Novo Assembly | SPAdes | 28 min 45 sec | 14.2 GB | 92% | Contigs: 142, Total length: 4,641,652 bp |
| Step 4: Assembly QC | QUAST | 4 min 38 sec | 1.9 GB | 68% | N50: 87,324 bp, L50: 18, GC%: 50.8% |
| **Total Pipeline Execution** | | **45 min 03 sec** | **14.2 GB** | **81% (avg)** | **Complete assembly with QC metrics** |

*Assembly quality metrics (N50: 87,324 bp, genome size: 4.64 Mbp) align with expected values for E. coli K-12 (reference genome: 4.64 Mbp). Test executed 5 times with <1% variation in metrics.*

#### Table 4: BioFrame Technology Stack

*Complete technology stack with version numbers and functional roles in the system architecture.*

| Layer | Technology | Version | Role/Purpose |
|---|---|---|---|
| Backend Framework | Python | 3.8+ | Core programming language |
| | Django | 4.2 | Web framework with MVT architecture |
| | Django ORM | 4.2 | Object-relational mapping |
| | Django REST Framework | 3.14 | RESTful API implementation |
| | Celery | 5.3 | Asynchronous task queue (optional) |
| Database | PostgreSQL | 15.3 | Relational database, metadata storage |
| | Redis | 7.0 | Caching and message broker |
| Containerization | Docker | 24.0.5 | Container runtime and image management |
| | Docker Compose | 2.20.2 | Multi-container orchestration |
| | Docker Python SDK | 6.1.3 | Programmatic container control |
| Frontend | HTML5 | - | Semantic markup |
| | CSS3 | - | Styling and responsive design |
| | JavaScript | ES6+ | Client-side interactivity |
| | Bootstrap | 5.3 | Responsive UI framework |
| | jQuery | 3.7 | DOM manipulation and AJAX |
| Visualization | Chart.js | 4.3 | Performance metrics visualization |
| | DataTables | 1.13 | Interactive data tables |
| Development Tools | Git | 2.40 | Version control |
| | pylint | 2.17 | Python code quality analysis |
| | coverage.py | 7.2 | Test coverage measurement |
| | Sphinx | 7.0 | Documentation generation |
| Operating System | Ubuntu Server | 22.04 LTS | Host operating system |

*All version numbers represent those used during development and testing. System is designed to work with compatible newer versions.*

#### Table 5: Concurrent Workflow Scalability Testing Results

*System behavior under varying concurrent workflow loads. Each workflow consisted of FastQC → Trimmomatic → FastQC (3-step pipeline).*

| Concurrent Workflows | Submission Latency | Avg. Completion Time | Peak Memory Usage | Peak CPU Usage | System Status |
|---|---|---|---|---|---|
| 1 (Baseline) | 0.8 sec | 12 min 15 sec | 6.2 GB | 78% | Optimal performance |
| 5 | 1.2 sec | 14 min 32 sec | 18.4 GB | 92% | Good performance |
| 10 | 1.8 sec | 18 min 45 sec | 28.6 GB | 98% | Acceptable performance |
| 20 | 3.4 sec | 28 min 18 sec | 30.8 GB (swapping) | 99% | Resource contention |
| 50 | 8.2 sec | 52 min 36 sec | 31.2 GB (heavy swapping) | 100% | Significant degradation |

*Test hardware: 8-core Intel Xeon, 32 GB RAM. Optimal performance maintained up to 10 concurrent workflows. Beyond this, memory pressure and CPU contention cause performance degradation. No system crashes or data corruption observed at any load level.*

#### Table 6: Software Testing Coverage by Module

*Code coverage percentages measured using coverage.py tool. Target threshold: 80% for core modules.*

| Module | Lines of Code | Lines Covered | Coverage % | Test Type |
|---|---|---|---|---|
| Workflow Orchestrator | 2,147 | 1,976 | 92% | Unit + Integration |
| Container Management | 1,523 | 1,387 | 91% | Unit + Integration |
| Logging System | 673 | 598 | 89% | Unit |
| Django Views | 1,842 | 1,544 | 84% | Unit + Integration |
| Database Models | 456 | 389 | 85% | Unit |
| API Endpoints | 892 | 758 | 85% | Integration |
| File Management | 634 | 521 | 82% | Unit |
| Utility Functions | 423 | 371 | 88% | Unit |
| **Total Core Modules** | **8,590** | **7,544** | **88%** | **Mixed** |

*Uncovered lines primarily include error handling for rare edge cases, debug logging statements, and deprecated code paths. All critical execution paths achieve >95% coverage.*

#### Table 7: Container Resource Requirements by Tool

*Memory and CPU requirements for each containerized bioinformatics tool based on typical genomic datasets.*

| Tool Name | Default Memory | Default CPUs | Image Size | Typical Runtime* |
|---|---|---|---|---|
| FastQC | 2 GB | 2 cores | 428 MB | 2-5 min |
| Trimmomatic | 4 GB | 4 cores | 156 MB | 5-15 min |
| SPAdes | 16 GB | 8 cores | 892 MB | 20-60 min |
| QUAST | 2 GB | 2 cores | 534 MB | 3-8 min |
| BWA | 8 GB | 8 cores | 312 MB | 15-45 min |
| SAMtools | 4 GB | 4 cores | 245 MB | 5-20 min |
| GATK | 8 GB | 4 cores | 1.2 GB | 20-90 min |
| MAFFT | 4 GB | 4 cores | 178 MB | 5-30 min |
| IQ-TREE | 8 GB | 8 cores | 234 MB | 10-120 min |
| MultiQC | 2 GB | 1 core | 456 MB | 1-3 min |

**Typical runtime based on E. coli-sized genome (~5 Mbp, ~100× coverage). Actual requirements vary based on genome size, sequencing depth, and analysis parameters. Memory and CPU allocations are configurable per workflow.*

#### Table 8: Database Schema - Core Entity Relationships

*Entity-relationship model for BioFrame platform implemented in PostgreSQL with Django ORM.*

| Entity | Description | Key Attributes | Relationships |
|---|---|---|---|
| **Tool** | Bioinformatics software package | name, category, description, icon | 1:N with Version |
| **Version** | Specific release of a tool | version_number, container_image, release_date | N:1 with Tool, 1:N with Parameter |
| **Parameter** | Tool input specification | name, type, required, default_value | N:1 with Version |
| **Workflow** | Sequence of tool executions | name, description, created_date, owner | N:N with Version (via steps), N:1 with User |
| **Run** | Execution instance of workflow | workflow_id, status, start_time, end_time | N:1 with Workflow, 1:N with Task |
| **Task** | Individual tool execution | run_id, tool_version_id, status, logs | N:1 with Run, N:1 with Version |
| **File** | Input/output data artifact | filename, path, checksum (SHA-256), size | N:N with Task (inputs/outputs) |
| **User** | Authenticated researcher | username, email, role, institution | 1:N with Workflow, 1:N with Run |

*Schema implements referential integrity with foreign key constraints. Status fields use ENUM types: pending, running, completed, failed, cancelled. All entities include timestamps (created_at, updated_at) for audit trails.*

#### Table 9: Usability Testing Results (n=5 participants)

*User acceptance testing results from graduate students in bioinformatics programs. Ratings on 5-point Likert scale (1=Strongly Disagree, 5=Strongly Agree).*

| Evaluation Criterion | Mean Score | Std. Dev. | Median | Interpretation |
|---|---|---|---|---|
| Ease of workflow creation | 4.2 | 0.4 | 4.0 | Easy to use |
| Clarity of user interface | 4.4 | 0.5 | 4.5 | Very clear |
| Real-time monitoring usefulness | 4.6 | 0.5 | 5.0 | Very useful |
| Error message helpfulness | 3.8 | 0.8 | 4.0 | Moderately helpful |
| Documentation quality | 4.0 | 0.7 | 4.0 | Good quality |
| Overall satisfaction | 4.0 | 0.6 | 4.0 | Satisfied |
| Likelihood to recommend | 4.4 | 0.5 | 4.0 | Highly likely |
| Preference vs command-line | 4.8 | 0.4 | 5.0 | Strong preference |

*Participants: 3 from University of Nizwa, 2 from collaborating institutions. Experience level: 1-4 years command-line bioinformatics. Testing conducted October 2024. Error message helpfulness rated lowest; improvements implemented based on feedback.*

#### Table 10: Comparison with Existing Workflow Systems

*Feature comparison between BioFrame and popular bioinformatics workflow management systems.*

| Feature | BioFrame | Galaxy | Nextflow | Snakemake |
|---|---|---|---|---|
| Web-based interface | ✓ | ✓ | ✗ | ✗ |
| Containerization support | ✓ (Docker) | ✓ (Docker/Singularity) | ✓ (Docker/Singularity) | ✓ (Docker/Singularity) |
| Linear pipeline execution | ✓ | ✓ | ✓ | ✓ |
| Parallel execution support | ✗ | ✓ | ✓ | ✓ |
| Real-time monitoring | ✓ | ✓ | Limited | Limited |
| Tool integration complexity | Low | Medium | Medium | Medium |
| Learning curve | Low | Low-Medium | High | Medium-High |
| Programming required | No | No | Yes (Groovy) | Yes (Python) |
| Multi-node deployment | ✗ | ✓ | ✓ | ✓ |
| Cloud platform support | Limited | ✓ | ✓ | ✓ |
| Data provenance tracking | ✓ | ✓ | ✓ | ✓ |
| Target user base | Lab researchers | Broad users | Bioinformaticians | Bioinformaticians |

*BioFrame optimized for ease of use and accessibility for laboratory researchers. Galaxy provides broader features but higher system complexity. Nextflow and Snakemake offer more advanced features but require programming expertise.*

---



Master's Thesis Research Project | Mohammed Al-Hinai

University of Nizwa | Department of Biological Science | Academic Year 2024-2025

All data tables are formatted for academic essay submission and can be printed to PDF or copied to Word documents.
